# Input-independent homeostasis of developing thalamocortical activity

**DOI:** 10.1101/2021.02.01.429056

**Authors:** Pouria Riyahi, Marnie A Phillips, Matthew T Colonnese

## Abstract

The isocortex of all mammals studied to date shows a progressive increase in the amount and continuity of background activity during early development. In humans the transition from a discontinuous (mostly silent, intermittently bursting) cortex to one that is continuously active is complete soon after birth and is a critical prognostic indicator in newborns. In the visual cortex of rodents this switch from discontinuous to continuous background activity occurs rapidly during the two days before eye-opening, driven by activity changes in relay thalamus. The factors that regulate the timing of continuity development, which enables mature visual processing, are unknown. Here we test the role of the retina, the primary input, in the development of continuous spontaneous activity in the visual system of mice using depth electrode recordings of cortical activity from enucleated mice *in vivo*. Bilateral enucleation at postnatal day (P)6, one week prior to the onset of continuous activity, acutely silences cortex, yet firing rates and early oscillations return to normal within two days and show a normal developmental trajectory through P12. Enucleated animals showed differences in silent period duration and continuity on P13 that resolved on P16, and an increase in low frequency power that did not. Our results show that the timing of cortical activity development is *not* determined by the major driving input to the system. Rather, homeostatic mechanisms in thalamocortex regulate firing rates and continuity even across periods of rapid maturation.

## INTRODUCTION

Spontaneous activity--sometimes called background or resting-state--is a pervasive feature of thalamocortical circuit function (Raichle 2010; Uddin 2020). The patterns of this activity are both a target and effector of arousal state modulation (McCormick et al. 2020). Multiple roles have been suggested for spontaneous activity: mediating attention, increasing signal-to-noise, resetting synaptic weights, and changing the functional connectivity of neurons (Harris and Thiele 2011; Froudarakis et al. 2019; Tononi and Cirelli 2020). Thus, it is not surprising that the acquisition of normal background activity is a key developmental checkpoint (Pavlidis et al. 2017). While phenomenologically well characterized in humans and animals (Tuge et al. 1960; Gramsbergen 1976; Vanhatalo and Kaila 2006; Andre et al. 2010), the circuit basis of background activity development is poorly understood (Wallois et al. 2020). This knowledge is crucial for treatment of early brain disorders and to improve the diagnostic utility of electroencephalography (EEG) for neonatal medicine (Pavlidis et al. 2017).

The background EEG of human infants fully resembles adult sleep-wake patterns a few months after birth, when sleep-spindles and large-amplitude slow-waves emerge (Tiriac and Blumberg 2016; Dereymaeker et al. 2017). Very early-born infants produce cortical activity that is highly discontinuous, with bursts of activity interrupted by long silent intervals of up to 60 seconds, which become shorter with age. The periods of activation consist of unique early patterns such as delta-brushes, *trancé alternate*, and temporal theta, the occurrence of which is both age- and region-dependent (Whitehead et al. 2017; Wallois et al. 2020). Neonatal injury, such as hypoxia/ischemia, can reverse gains in continuity (Stevenson et al. 2017), the recovery of which is a positive prognostic for neonatal brain function (van Rooij et al. 2005; Iyer et al. 2014).

All mammalian species yet examined show a similar developmental trajectory (Cirelli and Tononi 2015). The rodent visual system has proven to be a good model of cortical activity development, as the timing of critical events is well studied in both animals and humans, and control of the inputs can be easily achieved (Leighton and Lohmann 2016; Seabrook et al. 2017). In the primary visual cortex (V1) of rats and mice, spontaneous activity is discontinuous throughout most of the first two postnatal weeks, rapidly becoming continuous between P11 and P13, just before eye-opening (Colonnese et al. 2010; Shen and Colonnese 2016). This switch in the macro-patterning of background activity is contemporaneous with a change in the spectral patterns of activity as well as an increase in neural firing rates, reflecting an end to immature oscillations and the emergence of the adult-like ‘active’ state, as well as robust sleep-wake rhythms (Colonnese 2014).

Similar changes appear to occur in all thalamocortical pathways studied to date (Colonnese and Phillips 2018; Hanganu-Opatz et al. 2021), but the mechanisms are not understood. Recent recordings in the visual relay thalamus, the dorsal lateral geniculate nucleus (dLGN), suggest that changes there cause the maturation of the background EEG in V1 (Murata and Colonnese 2018). However, activity of the major input to dLGN, retinal ganglion cells (RGCs), also becomes more continuous around eye-opening (Demas et al. 2003). Thus, it is possible that the acquisition of adult-like background activity in dLGN and V1 simply reflects retinal maturation.

Here we examine the role of retina in timing the development of background activity in V1 by removing both eyes during the early period of discontinuity and recording V1 activity development through the eye-opening period. While we find subtle differences in the patterns of early activity in enucleated animals, the gross acquisition of continuous background activity is very similar in enucleated and control animals. We conclude that powerful homeostatic mechanisms within thalamocortex control activity development in V1, and that a genetically programmed shift in the thalamus (or its modulatory inputs) controls the timing of cortical background activity development, without instruction from the retina.

## Materials and Methods

### Animal care

Animal care and procedures were in accordance with *The Guide for the Care and Use of Laboratory Animals* (NIH) and approved by the Institutional Animal Care and Use Committee at The George Washington University. Postnatal day (P)0 is the day of birth. C57BL/6 were obtained from Hilltop Lab Animals (Scottsdale, PA) as timed pregnant females, and kept in a designated, temperature and humidity-controlled room on a 12/12 light/dark cycle and examined once per day for pups. For bilateral enucleations, carprofen (20 mg/kg) in saline was injected 1 hour prior to surgery to reduce pain and inflammation. Surgical anesthesia was induced with 3% isoflurane vaporized in 100% O_2_, verified by tail-pinch. An incision was made in the eyelid (P6) and the globe of the eye was removed using forceps. The eye socket was filled with sterile surgical foam (GelFoam) and the eyelid closed using a tissue adhesive (Vetbond). Pups were post-operatively monitored and received follow-up injections of carprofen daily for 2 days. Sham control animals received identical treatment, including eyelid puncture with the tip of a suture needle, without enucleation or GelFoam.

### In vivo electrophysiology

Procedures were as previously described (Shen and Colonnese 2016). Carprofen (20 mg/kg) in saline was injected 1 hour prior to surgery to reduce pain and inflammation. Surgical anesthesia was induced with 3% isoflurane vaporized in 100% O_2_, verified by toe-pinch, then reduced to 1.5-3% as needed by monitoring breathing rate and toe pinch response. An electrical heating pad (36°C) provided thermoreplacement. For attachment of the head-fixation apparatus, the scalp was excised to expose the skull, neck muscles were detached from the occipital bone, and the membranes were removed from the surface of the skull. Topical analgesic was applied to the incision of animals older than P8 (2.5% lidocaine/prilocaine mix, Hi-Tech Pharmacy Co., Inc., Amityville NY). Application to younger animals was lethal. The head-fixation apparatus was attached to the skull with grip cement (Dentsply, Milford DE) over Vetbond™ tissue adhesive (3M). Fixation bar consisted of a custom manufactured rectangular aluminum plate with a central hole for access to the skull. After placement, the animal was maintained with 0.5-1% isoflurane until the dental cement cured, after which it was allowed to recover on the warmed table.

For recording, animals were head-fixed via the plate and body movements were restricted by placement in a padded tube. Body temperature was monitored via thermometer placed under the abdomen, and maintained above 33C via thermocoupled heating pad (FHC, Bowden ME). Body motion was monitored with a piezoelectric device placed below the restraint tube. For electrode access, a craniotomy was performed thinning the skull if necessary and resecting small bone flaps, to produce a small opening (∼150-300 µm diameter). Primary visual cortex targets were determined by regression of adult brain lambda-bregma distances: 1.5-2.5mm lateral and 0.0-0.5 mm rostral to lambda. All recordings were made using a single shank, 32 channel array arranged in two parallel lines of contacts (A1×32-Poly2-5mm-50s-177, NeuroNexus Technologies, Ann Arbor MI). The electrode penetrated the brain orthogonally to the surface and was advanced to a depth of 500-800µm using stereotaxic micromanipulator until the top channels touched the cerebral-spinal fluid. Isoflurane was withdrawn and the animal acclimated to the setup for at least 60 min prior to recording. Unless otherwise noted recordings were 30min. All recording was performed in the dark (<0.01 Lumens). Following recordings, all animals were sacrificed by anesthetic overdose followed by decapitation.

### Data acquisition and analysis

Data was amplified 192V/V and digitized at 30 kHz on SmartLink Headstage and recorded with the SmartBox 1.0 (Neuronexus, Ann Arbor MI). Recordings were imported in Matlab using Neuronexus supplied macros and custom code. Depth (d)EEG signals were derived by down-sampling wide-band signal to 1 kHz after application of 0.1-350Hz zero-phase low-pass filter applied. To remove common mode noise and volume conduction from cortical structures, signals were referenced to a sub-cortical contact in layer 6 with minimal spiking. For multi-unit activity, channels with high noise (outside the brain or bad contacts) were manually selected and eliminated and the remaining channels saved as binary structures for spike sorting using Kilosort (Pachitariu et al. 2016). To derive ‘cleaned’ multi-unit activity, clusters were identified using the default parameters with the following exceptions: ops.Th = [3 6 6]. Noise clusters were manually removed using the spike sorting GUI Phy (Rossant et al. 2016) and all remaining clusters were collapsed by central contact as determined by the mean spike-waveform minima. Spike-times were binned at 1 ms.

All analyses were performed in Matlab. Recordings were not included in the analysis if they contained spreading depression (n=4) or if the merged L2-4 or L5-6 MUA was below 0.1Hz (n=7). One additional animal was eliminated because the file was corrupt and could not be imported. Before analysis, periods of movement as identified by the piezo signal, were removed. Channels were divided into superficial (L2-4) and deep (L5-6). Superficial layers were identified by depth and the presence of high-frequency (>10 Hz) power (Colonnese and Khazipov 2010). Spike rates and continuity were derived from the 4 channels in each (L2-4 & L5-6) with the highest mean spike rate. For spectral analysis, the re-referenced signal from a contact in the center of L2-4 was selected. Spectral decomposition of the dEEG signal used the multi-taper method (Mitra and Bokil 2007). Spectra were calculated for 1s windows (time-bandwidth product 3 and number of tapers 5). Window width and time-bandwidth products were chosen empirically to maximize the spindle-burst frequencies in young animals. Normalized mean power for each animal was calculated by averaging all windows during non-movement periods then dividing by mean 1-60 Hz power. Division into active and inactive periods followed loosely the method of (Renart et al. 2010) as implemented previously in neonatal mice (Colonnese et al. 2017). The smoothed multi-unit spike rates were calculated by summing spike times from 4 highest responding channels in superficial or deep layers and applying a gaussian window with 25ms half-width (Fig 3A). Based on this smoothed spike-rate vector, active and inactive periods were identified with a threshold set to 10% of maximal spike rate for that mouse and layer. Inactive periods of less than 100ms were folded into the adjacent active periods. For a subset of analyses (Fig 4C) the gaussian half-width was increased to 200ms. For analysis of active and silent period distributions, the occurrence of each duration was accumulated into one of 50bins, with log distributed widths, between 10^1^ and 10^5^ ms, and normalized to the total occurrences. To quantify the change in distributions between days as a single variable, we calculated the absolute difference between the normalized distributions for each animal in each group against each of the animals in the reference group(P16-17). For each group and age these pair-wise differences were averaged.

### Statistical Procedures

All data are mean +/-standard deviation, except for the distribution differences which are 95% confidence intervals. Statistical test applied is noted along with results. Statistical tests were applied in Matlab using the inbuilt functions (‘ttest2’, ‘anova’). Significance statements regarding normalized frequency power distributions were calculated by permutation analysis following the permutation test method of Cohen (Cohen 2014), using custom macros. Frequency resolution was 1 Hz from 1-60Hz. The significance threshold was p<0.05.

## RESULTS

Early enucleations were made at P6, a developmental time when enucleation does not produce gross sprouting of other sensory inputs to dLGN or cause reorganization of cortico-cortical connections (Olavarria and Hiroi 2003). We first determined the acute effects of early bilateral enucleation on spontaneous activity in the visual cortex of mice. Recordings were made in five P6 mice (2 liters) using a single-shank 32-channel multi-electrode array inserted in the monocular zone of V1. After baseline recordings, the mice were anesthetized and binocular enucleation performed with the electrode in place (Fig. 1A).

**Figure 1.**
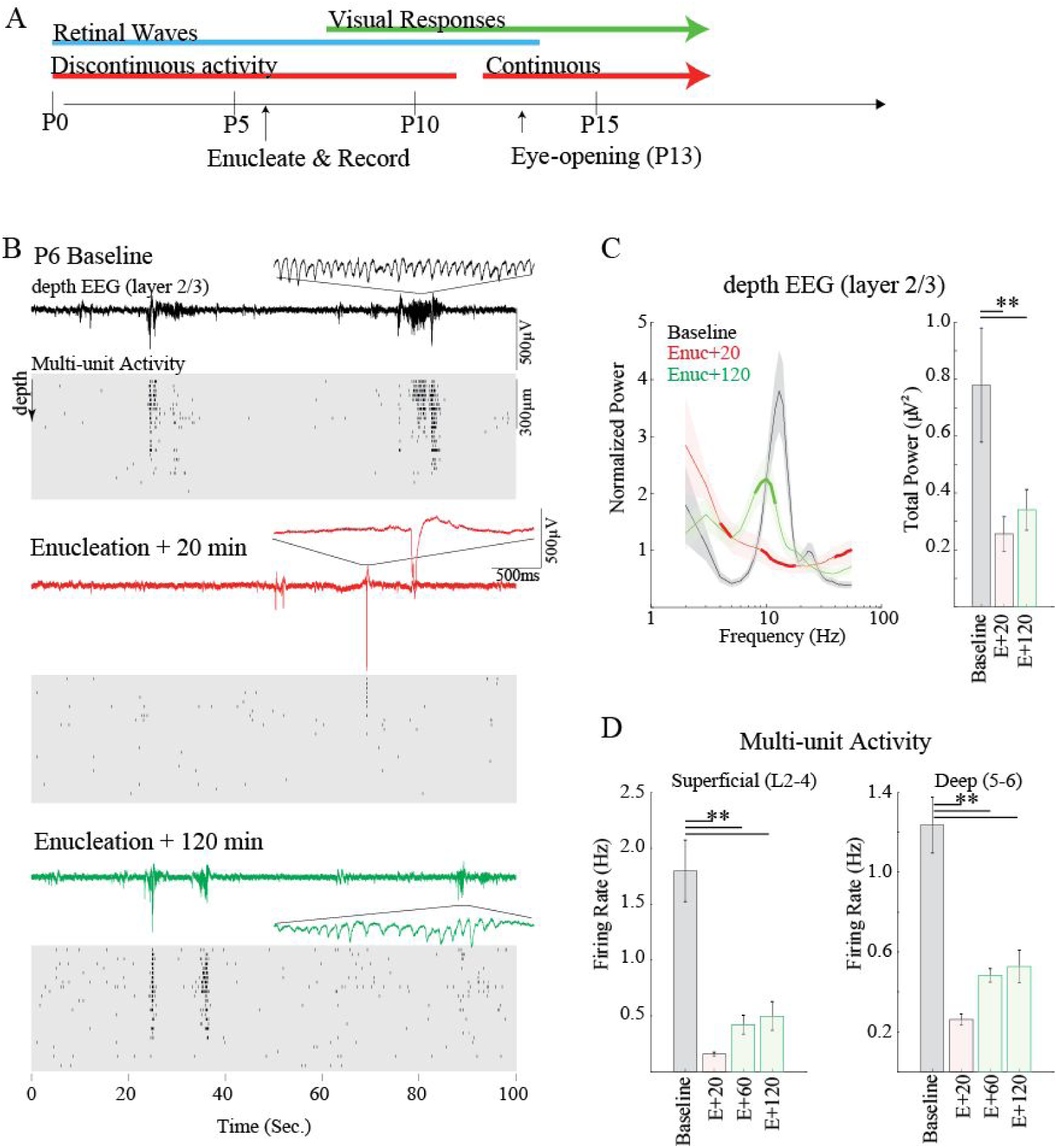
Enucleation acutely reduces activity and spindle-bursts in mouse visual cortex. **A**. Overview of major developmental stages in visual cortex of the mouse relative to acute enucleation on the 6^th^ post-natal day (P6). **B**. Representative example of spontaneous activity before (Baseline), 20 and 120 minutes following bilateral enucleation. Top line shows depth EEG, with detail of one spindle-burst. Cleaned multi-unit raster of spike times is below. Each line is from a single contact with 25µm of depth between contacts. Note rapid loss of rapid oscillations in bursts as well as spontaneous spiking at Enuc+20min, with some recovery of both by Enuc+120 min. **C**. Population mean (n=5) spectral analysis for dEEG. Left: mean ± StDev spectral power distribution. Thick line segment shows regions of significant difference from preceding condition (i.e. Baseline for Enuc+20, and Enuc+20 for Enuc+120) determined by premutation analysis. Right: Summed 1-60Hz power. Animals lose and then recover spindle-burst frequencies following enucleation, but total power remains low. **D**. Population mean multi-unit firing rates. ** = p<0.01 by Tukey post-hoc test.

As previously described (Shen and Colonnese 2016), baseline spontaneous activity on P6 was largely restricted to regularly repeating 1-10s periods of high-firing rates and prominent 10-20Hz oscillations in the superficial layers (Fig. 1B & C). These oscillatory periods, called slow-activity transients (Colonnese and Khazipov 2010), consist of multiple spindle-bursts which are the result of spontaneous retinal wave input to the thalamus (Murata and Colonnese 2016), which is then conveyed to topographically appropriate locations in visual cortex (Ackman and Crair 2014; Leighton and Lohmann 2016; Kummer et al. 2016). After enucleation, spontaneous cortical activity was recorded for two hours (Fig. 1B) and analyzed in 20 minute windows beginning 20min, 60 min and 120 min after enucleation (Table 1). Immediately after recovery from anesthesia (E+20min), spindle-bursts were absent as evidenced by an elimination of the prominent 10-18 Hz bump in the normalized frequency power of the layer 2/3 depth (d)EEG (local field potential)(Fig. 1C). Spontaneous activity overall was severely curtailed as well, as evidenced by a 3-fold drop in 1-60Hz power (Table 1). Likewise, the multi-unit firing rate dropped by 90% in superficial layers and 79% in deep layers. The remaining activity consisted of isolated firing and short (<1s) bursts. Such a dramatic loss of activity is similar to that observed following retinal activity blockade in rats by microinjection at similar ages (Murata and Colonnese 2016), suggesting that it is due to the loss of retinal ganglion cell activity, not unexpected side-effects of retinal trauma. Total spontaneous activity remained suppressed at 60 and 120 minutes post enucleation. By 120 minutes, however, the cortex had begun to produce spindle-burst oscillations (Fig. 1B), and as a result, we observed a significant increase in normalized frequency power 8-15Hz (Fig. 1C), though the total 1-60Hz power remained similar to 20 minutes post-enucleation. Mean firing rates tripled between 20 and 120 min post-enucleation in superficial layers and doubled in deep layers (Fig. 1E), though this increase was not significant by post-hoc test, likely because of the total variability introduced by pre-enucleation firing rates.

**Table 1.**
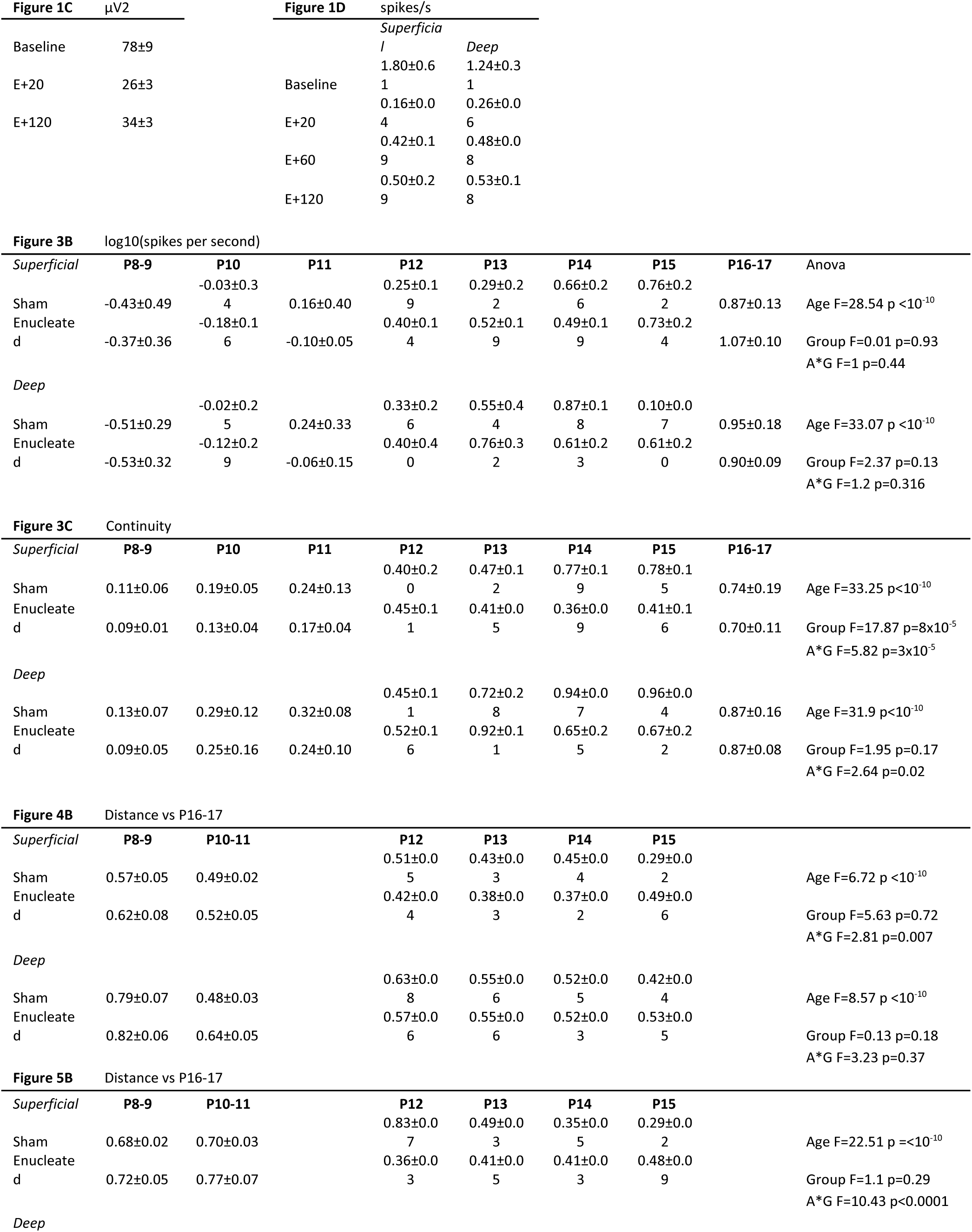
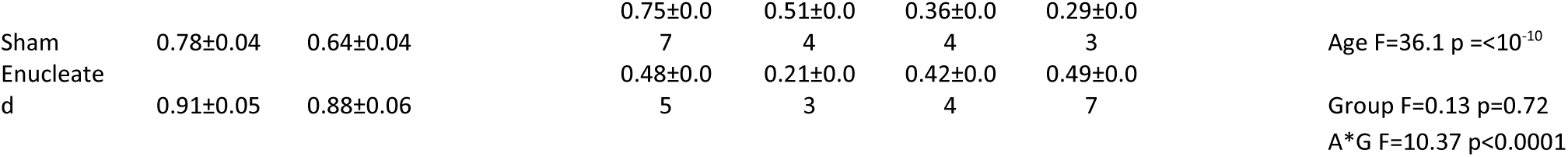
Mean and Standard deviations. Two way ANOVA results shown for the group comparisons.

Together these acute experiments show that enucleation at P6 has a profound effect on the activity of visual cortex. However, the retina is not the only input capable of driving V1 activity at this age, and plasticity to restore normal activity patterns begins rapidly.

### Chronic effects of P6 enucleation

To determine the degree to which enucleation changes the developmental trajectory of spontaneous activity we performed bilateral enucleation or sham surgery on littermates of P6 mice. Spontaneous activity was then acutely recorded from enucleated and sham control littermates at single day intervals between P8 and P17 (Fig. 2). Previous work (Shen and Colonnese 2016) indicated that P12-15 is a time of rapid change in the activity of visual cortex; Thus, for quantification we analyzed the following age groups: P8-9 (n=5 Sham & 5 Enucleated), P10-11 (n=7 & 7), P12 (n=4 & 5), P13 (n=4 & 4), P14 (n=5 & 9), P15 (n=5 & 5), and P16-17 (n=4 & 3).

**Figure 2.**
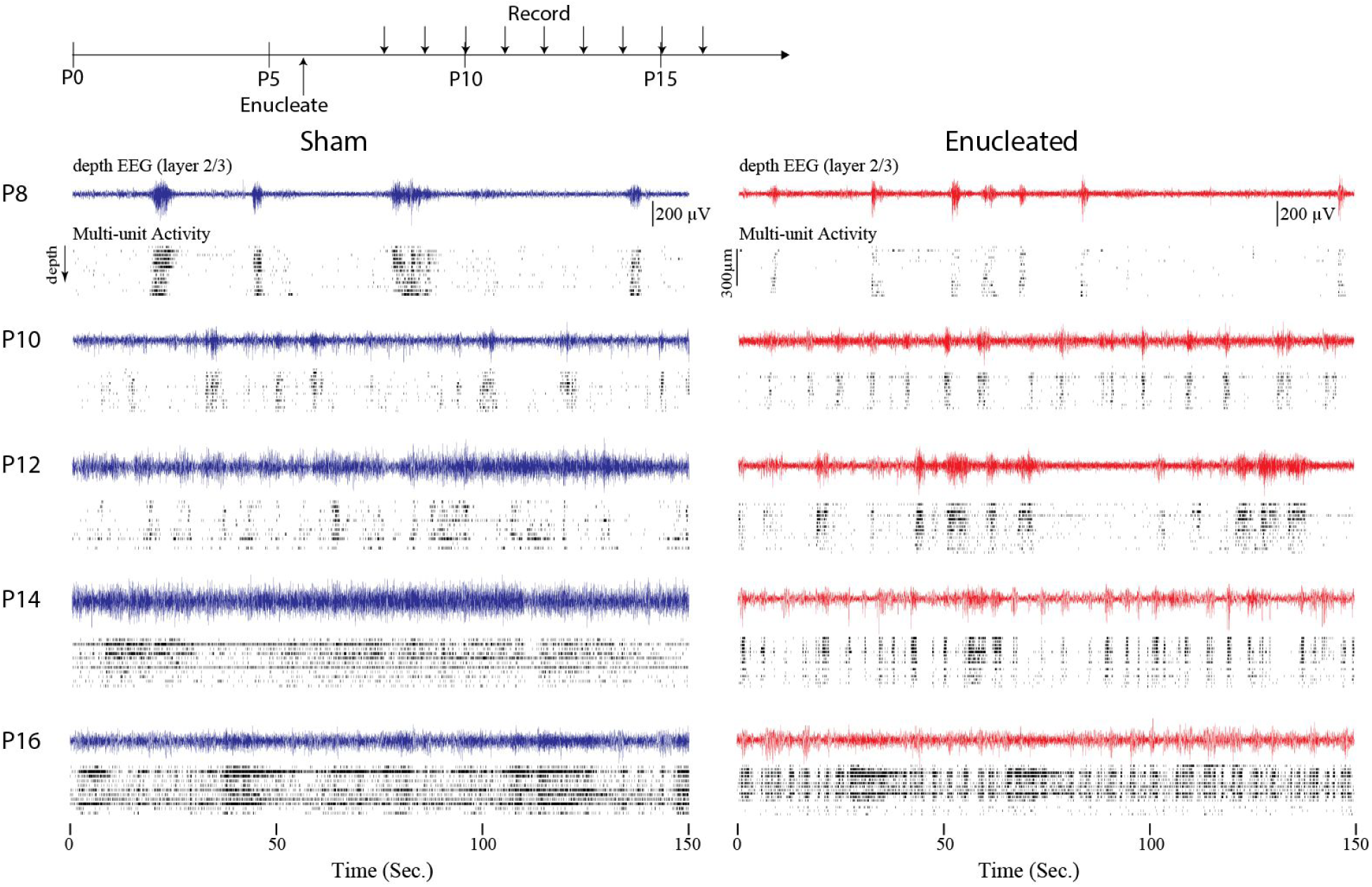
Developmental trajectory of cortical activity following bilateral enucleation. Top: Timing of enucleation and recordings. Recordings are acute and from different animals each day. Below: depth EEG and cleaned multi-unit rasters throughout the depth of visual cortex in two-day intervals. Enucleated animals have visibly different patterns of activity vs. silence but total activity develops on a similar trajectory, and those enucleated resemble sham control animals by P16.

Qualitatively, spontaneous activity developed similarly in enucleated and sham control animals (Fig. 2). Between P8 and P11 the visual cortex of both groups displayed long periods of silence interrupted by multi-second bursts of rhythmic (8-20Hz) activity. The bursts of rhythmic activity appeared shorter and more regular in enucleated animals, particularly at P8 and P10. After P12 the continuity of the spontaneous activity increased in both groups. By P16 significant (>1s) silent periods were absent in both groups. The detailed trajectory of this development varied between sham and enucleated, however. Between P12 and P15 the activity of enucleated animals consisted of many short bursts of activity, while sham animals showed longer periods of continuous activity, particularly around P14-15, which was the only age at which the two groups could be reliably distinguished by eye.

We first quantified multi-unit firing rates in superficial (L2-4) and deep (L5-6) layers (Fig 2A). The developmental trajectory of MUA firing rates in superficial or deep layers was not significantly different between treatment groups (Fig. 2B): Two-way ANOVA found no effect of group, a significant effect of age (x) and no interaction between the two (Table 1); Furthermore, post-hoc analysis revealed no differences between groups for any age group. For both groups, the first day that mean firing rates were significantly different from those observed on P8 was P12. No significant differences between P12 and older groups were observed for either group. Thus, the developmental trajectory of firing rates is independent of retinal presence, even though during normal development retina drives around 80-90% of cortical activity during the first two post-natal weeks (Murata and Colonnese 2016).

To assay the development of temporal patterning, particularly the continuity of activity, we used multiple measures based on separating silent and active periods (Fig 2A). To separate these periods a gaussian filter (20ms half-width) was applied to the summed superficial or deep layer MUA and a threshold (3% peak) applied to define silent and active periods (Colonnese et al. 2017).

Continuity was measured as the proportion of total time spent in an active period. Unlike mean firing rates, the developmental trajectory of continuity revealed a complex interaction between retina-dependent and independent influences, with retina influencing superficial layers more than deep layers (Fig. 3C). Two-way ANOVA on superficial layer continuity revealed a significant effect of age and group and an interaction between the two (Table 1). This was also observed for deep layer continuity. For superficial layers, post-hoc analysis revealed that the first age at which continuity was significantly increased over P8 was P12 for both treatment groups; for deep layers it was P12 for control and P13 for enucleated. Sham animals showed a second significant increase in continuity between P12 and P14 in both layers. Enucleated animals, however, did not express this second increase in continuity until P16-17. As a result, superficial layer continuity was significantly different between sham and enucleated animals on both P14 and P15. In deep layers, continuity was reduced at these ages, but not significantly relative to control.

**Figure 3.**
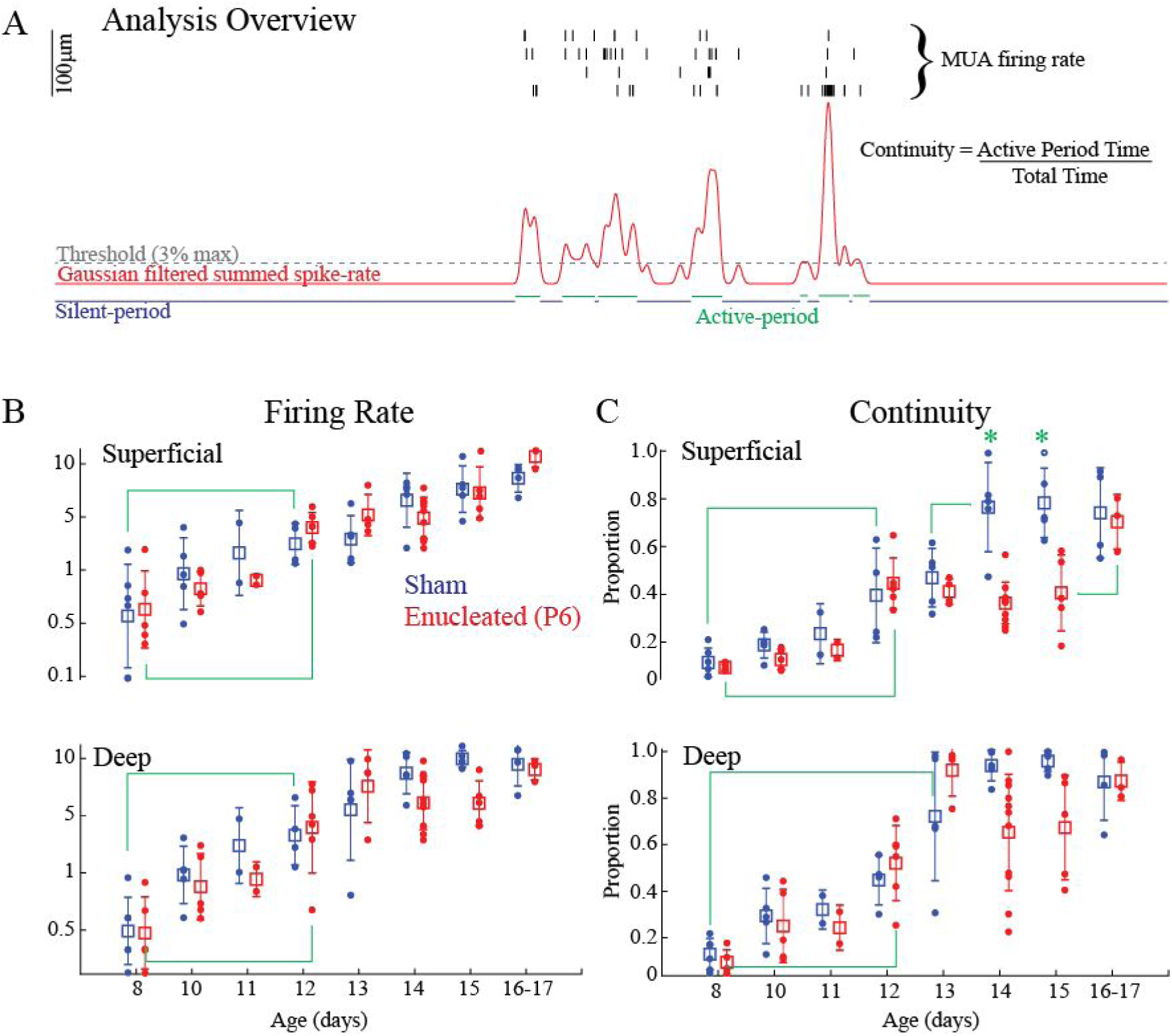
Development of activity in visual cortex is largely independent of retina. **A**. Schematic of analysis for Fig. 3-5 & 7. Four channels with maximal multi-unit firing are selected for superficial (L2-4) and deep (L5-6) layers and spike times summed. Recordings are divided into silent and active periods based on a threshold of smoothed firing rate vector for each layer. A general measure of continuity reflects the total amount of time spent in the active ‘state’. **B**. Developmental trajectory of multi-unit firing rates by age. Mean and StDev are shown by box and whiskers, with individual animals as dots. Green line shows the first days with significant difference from P8 by Tukey post-hoc test (p<0.05). No older ages were significantly different from P12. **C**. Developmental trajectory of continuity. Green lines show closest within-group significant differences for each age. Green asterisks show differences between groups for that age (p<0.05). Enucleated mice show an initial increase in continuity at P12, but a second jump at P14 is delayed by two days.

Thus continuity develops in two stages. Continuity first increases around P12, even in the absence of the retina. A second increase occurs around P14, and this second increase is slightly delayed following long term enucleation in superficial layers.

### Development of active and silent periods

While the bulk measures of firing-rate and continuity showed that development of background activity in the visual cortex is largely independent of retina, visual inspection of the animals suggested that the micropatterning of activity and silences are at least partially retinal dependent (Fig.2). To capture this difference, we examined the duration of active and silent periods. We first measured the duration of active periods, as defined by the summed MUA activity across superficial or deep layers using a 20ms gaussian filter for smoothing (Fig 3A). Surprisingly given the obvious differences in pattern and continuity, the distribution of active period durations did not change significantly across development either in control or enucleated animals (Fig. 4A). To analyze this statistically, the absolute difference between distributions was measured, with P16-17 animals serving as the reference (Fig. 4B). This analysis indicated constant difference across all ages for both superficial layers and deep layers, with no effect of age or group (Table 1). This statistical lack of development of active period duration was hard to reconcile with visual inspection of the data in young animals, which clearly showed long duration events that were not present in the enucleated group. Because the slow activity transients, driven by retinal waves at these ages, consist of multiple clusters of shorter events, we hypothesized that a wider filter would allow for the capture of these longer events. Therefore, we calculated the activity period duration distributions using a 200ms half-width Gaussian filter and minimum silent period of 500ms at P8-9 and P10-11 (this analysis doesn’t work in older, continuous animals). This analysis revealed a reduced occurrence of long duration (>∼3 seconds) events in superficial and deep layers at P8-9 and in deep layers at P10-11 (Fig. 4C). Thus, while the temporal structure of activity ‘bursts’ is largely unchanged following enucleation, long-duration activity periods driven by spontaneous retinal waves are absent when the eyes are removed.

**Figure 4.**
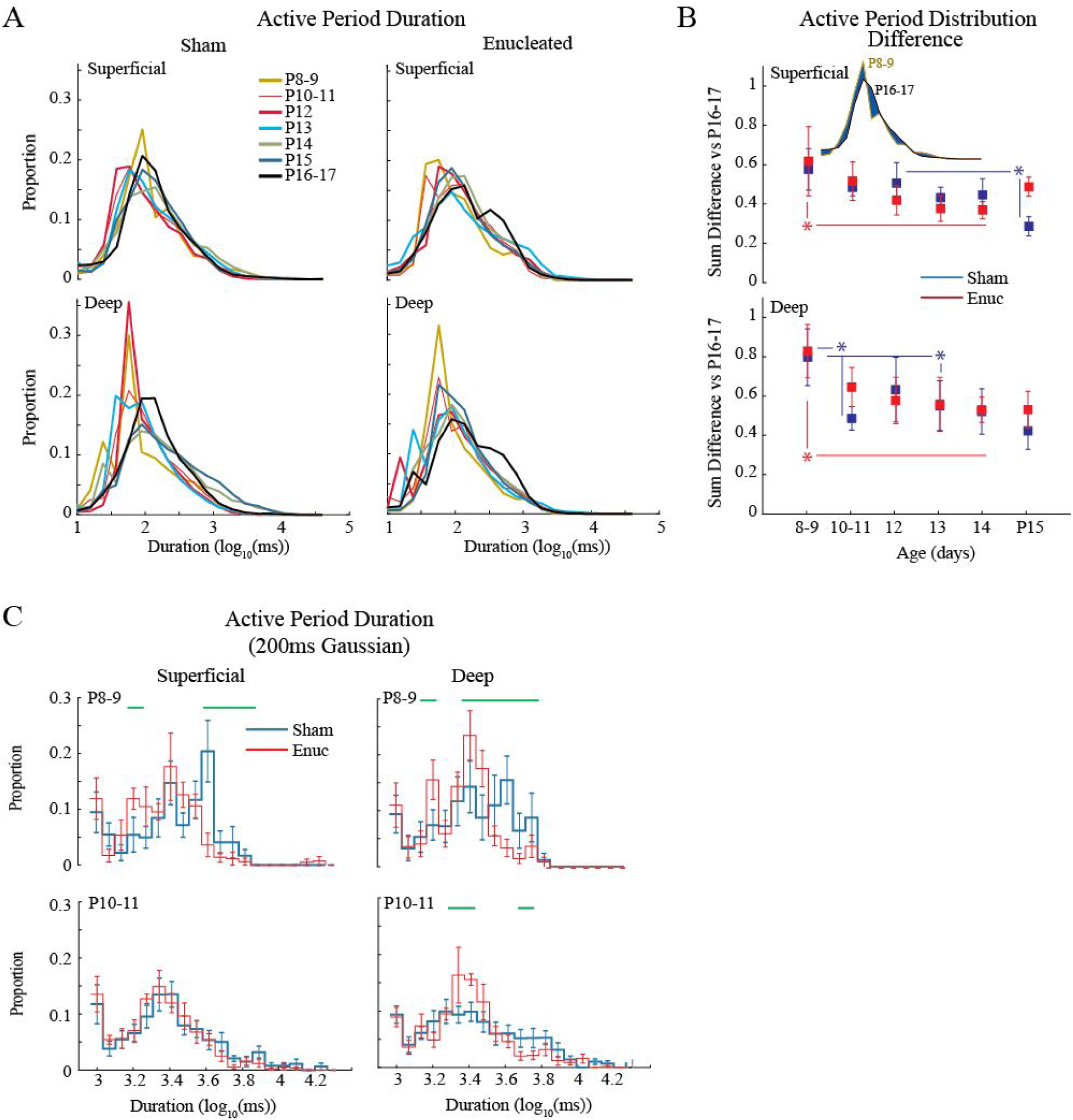
Development of active periods. **A**. Mean distribution of active period duration by age group. Active period duration is largely similar between ages and treatment groups. **B**. Quantification of active period duration development. Absolute distance between distributions (shown in inset as blue area) is calculated for all animals in the target age against the oldest (p16-17) group. The mean and 95% C.I. for these distances is plotted by age. Results show a steady increase in similarity to P16-17 distributions for both treatment groups. Only P15 animals (superficial layers) showed a difference between sham control and enucleated. **C**. Population mean of active period duration distributions using a longer smoothing filter to eliminate short silent periods. Green lines indicated durations with a significant difference in proportion as determined by permutation analysis (p<0.05).

In contrast to active periods, silent period duration was strongly predictive of age in both control and enucleated animals (Fig. 5A) as long silences were largely eliminated by P16-17 in all groups and layers. The absolute distance between silent period distributions for each age (vs. P16-17) showed significant effects for age and group, and an interaction between the two (Table 1). Post-hoc test revealed that silent period distributions rapidly become more like P16-17 between P12 and P14 in control animals, both in the superficial as well as deep layers (Fig. 5B). In enucleated animals this maturation occurred a day earlier, between P10-11 and P12. Because of the extensive changes in silent periods durations, we more closely examined the differences between control and enucleated groups for each age range (Fig. 5C). These comparisons show that on P12 the silent periods in enucleated animals become significantly shorter than sham. By P14 however, short periods of silence are dominant in control animals, while enucleated animals developed a significant subset of silent periods longer than one second. By P16-17 enucleated animals eliminated these long silent periods and resembled sham animals.

**Figure 5.**
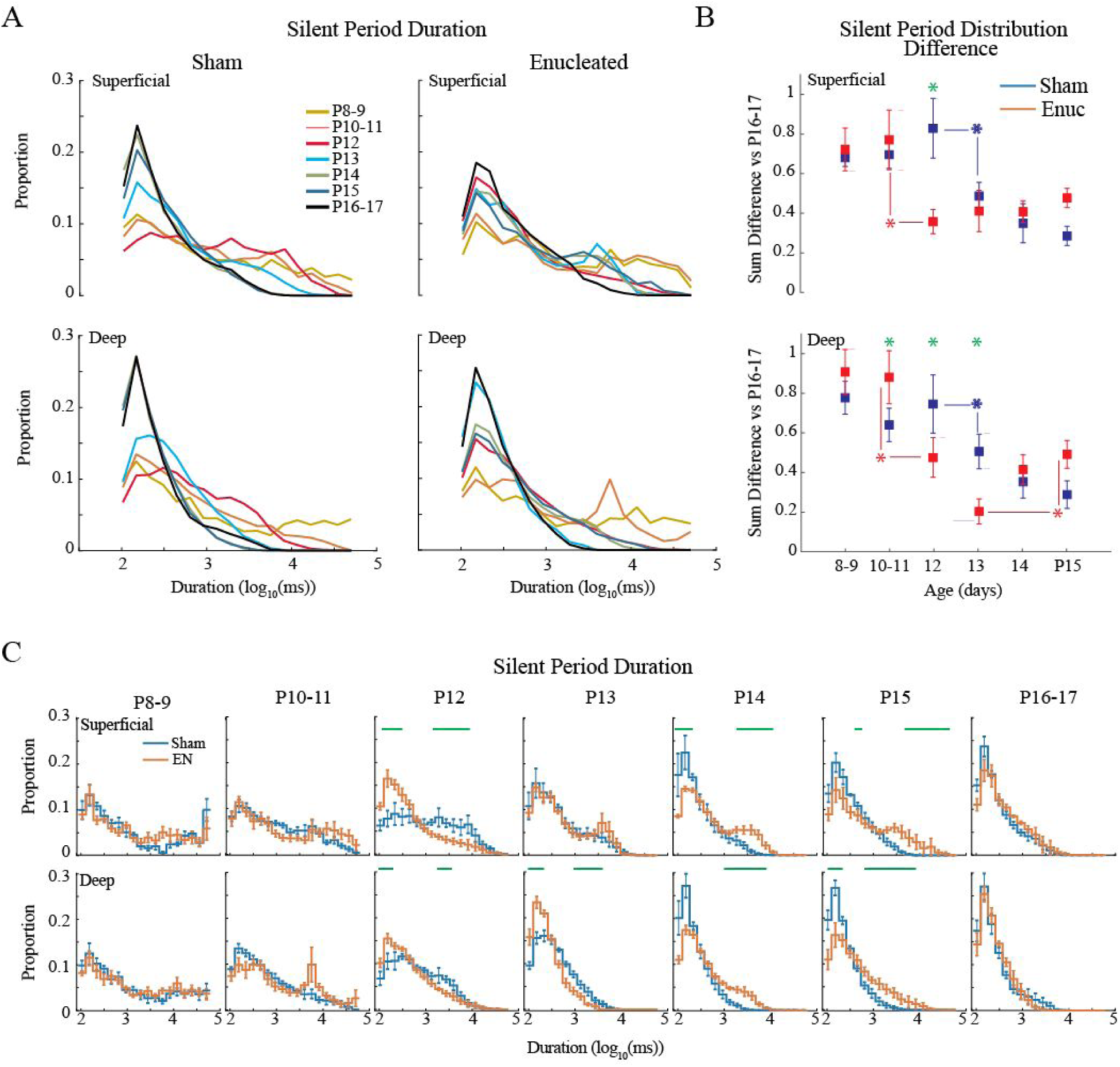
Development of silent periods. **A**. Population means distributions of silent period duration. Both groups and layers show a robust shift from long duration silent periods to short silent periods over the second and third post-natal weeks. **B**. Mean ± StDev of absolute distribution distance from P16-17 for each age. Green lines indicate significant difference p<0.05) by Tukey post-hoc test for age-adjacent groups. Green asterisks indicate significant differences between groups at a given age. **C**. Population means ± StDev of silent period duration distribution by age group. Green lines denote durations with significant differences (permutation analysis) between the groups. Enucleated animals show an increase in short duration silent periods one day early (P12 not P13) but then fail to further decrease long duration periods at P14 as in sham controls. Both groups have eliminated long duration periods by P16-17.

Together our results show that the basic trajectory of background activity, particularly the developmental reduction in network silence, is not a result of retinal activity and its timing is not critically dependent on retina. However, retinal input does appear to influence the specific timing of certain events, including shortening of silent period duration on P14 which leads to a delay in the development of normal continuous background activity until P16-17 in enucleated animals.

### Role of retina in neural oscillation development

The normal development of background activity includes a transformation of the micro patterning of activity, as observed in the frequency power of the dEEG. During the first eleven days, activity in superficial layers is dominated by spindle-burst oscillations, whose characteristic frequency increases with age. These spindle-bursts develop into the broad-band beta-gamma activity observed during visual activation and wakefulness. Slow-wave activity during quiescence is first observed at P10 and becomes prominent by P12 (Shen and Colonnese 2016). To see if this developmental trajectory is dependent on the retina, we examined the mean power of the layer 2-3 dEEG in sham and enucleated animals (Fig. 6). At P8, active periods included prominent spindle-burst oscillations in both groups. However, spectral power was higher in the enucleated group at 3-11Hz. Mean spectral power for enucleated animals peaked near 10 Hz rather than the 20 Hz seen in sham animals (Fig. 6B). Normal animals increase the frequency of spindle-burst oscillations from ∼10 Hz at P6 to 20 Hz at P8 (Shen and Colonnese 2016), suggesting that enucleation froze development of the oscillator at the time of enucleation. In P10-11 animals the groups showed significantly elevated power between 4 and 10 Hz, and reduced power at 20-60 Hz, suggesting a continued delay in the development of the central generator of spindle bursts. With the switch to the more mature patterns of activity at P12, significant differences in normalized power disappeared. Starting at P14 through all the ages examined in the study, enucleated animals had significantly elevated power at low frequencies and reduced power at high frequencies.

**Figure 6.**
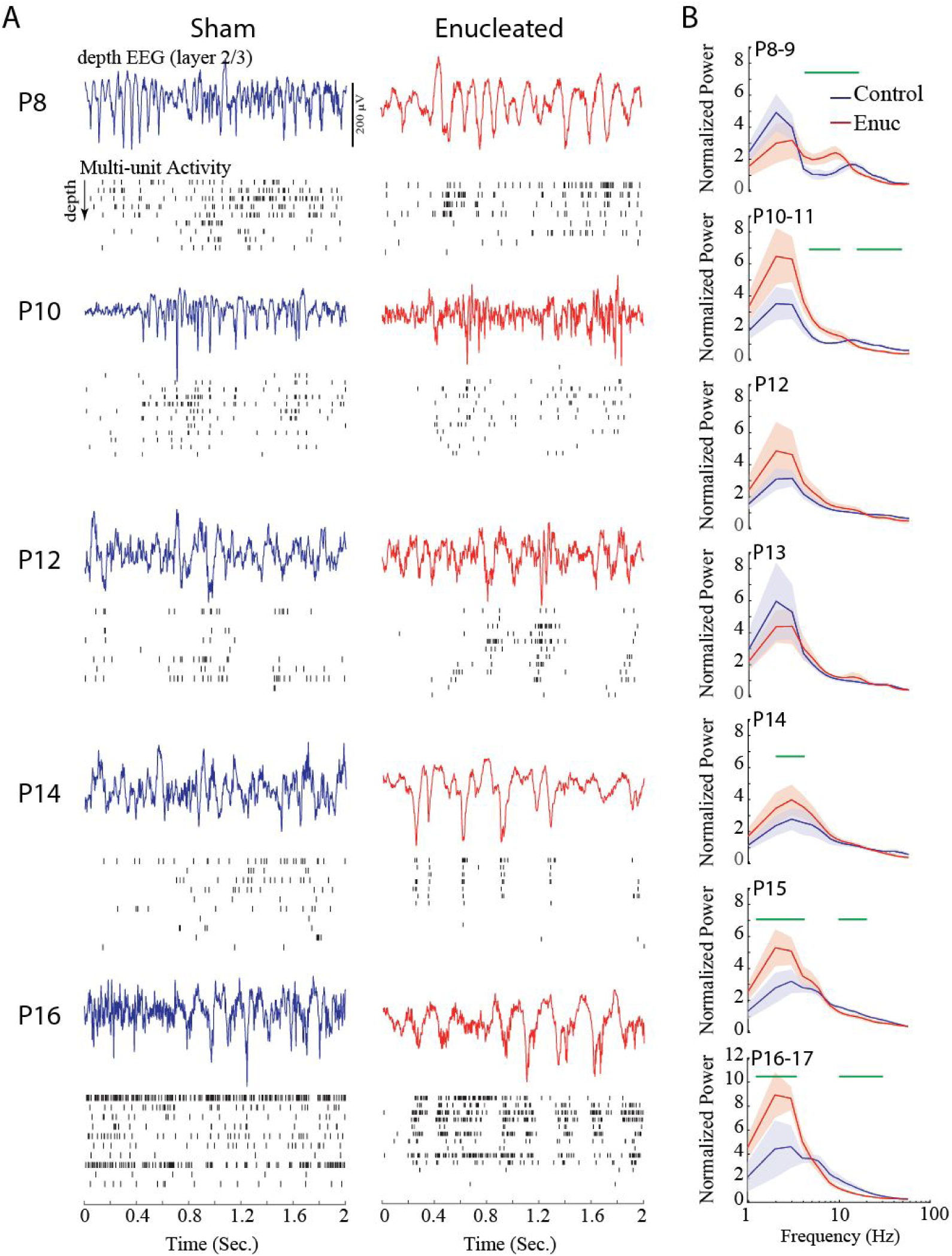
Role of retina in the development of micro-temporal patterning. **A**. Representative dEEG during active periods and multi-unit spike rasters through the depth of cortex. Activity periods were chosen to best represent spectral differences. **B**. Mean ± StDev spectral power distributions by age group. Green lines show frequencies with significant (p<0.05) differences between groups calculated by permutation analysis. Enucleated animals show reduced spindle-burst frequency at P8 and increased power in slow frequencies beginning at P14.

Together our analysis of dEEG frequency power confirms our earlier results that the fundamental developmental trajectory of background activity in visual cortex, specifically the switch from spindle-burst oscillations to counterbalanced low and high-frequency oscillations, occurs normally in enucleated animals. Enucleation, however, does result in a mild/moderate delay and some long lasting changes in how these early and late activity patterns are expressed.

### Late enucleation and activity development

Our P6 enucleations revealed a two-stage development of continuity, with an initial step occurring on P12 and a second on P14. Eye-opening in our mice naturally began on P13, suggesting the hypothesis that pattern vision drives the second step. Furthermore, it is possible that the establishment of normal background activity has a critical period which begins only after the end of the early, discontinuous, activity period. To test these hypotheses we performed enucleations and sham enucleations on littermates at P13, after the first switch and before eye-opening. Sham control animals also had their eyes opened during sham surgery under the microscope used for enucleations. Animals were recorded two days later at P15 when continuity, spectral and silent duration differences were most pronounced following P6 enucleation. In contrast to the early enucleations, P13 enucleation had no discernible effect on background activity at P15 (Fig. 7). Multi-unit firing rates, continuity, silent period duration and normalized frequency power all showed no significant differences between sham and enucleated groups. These data show that the onset of visual experience *per se* is not the driver for the second step in activity development. They further suggest that it is not the absence of retina *per se* that causes the differences in activity at P14-15.

**Figure 7.**
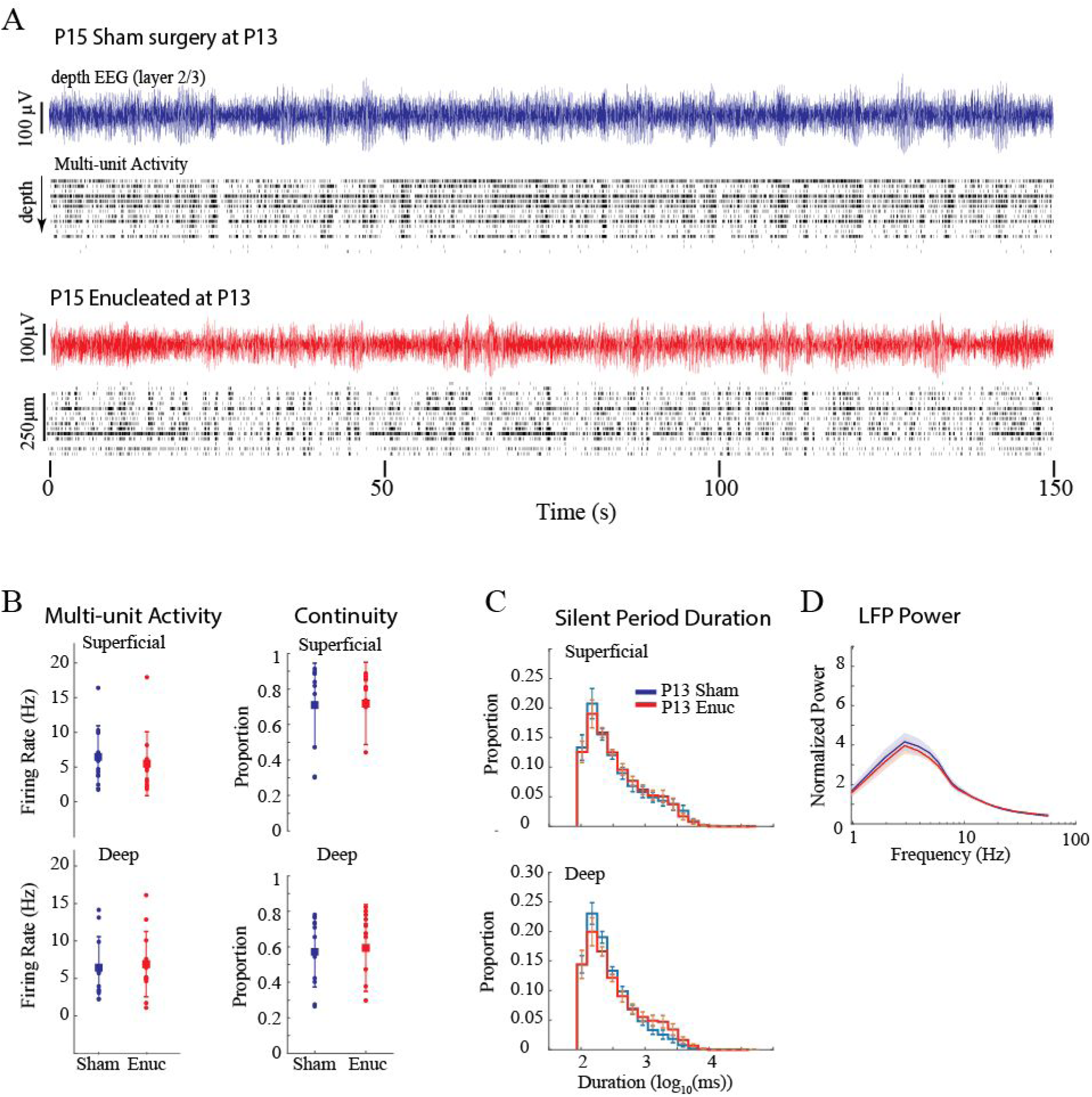
P13 Enucleation does not alter activity at P15. **A**. Representative example of spontaneous activity for sham control and enucleated animals. **B**. Population means ± StDev for firing-rates and continuity. **C**. Population means ± StDev for silent period duration. **D**. Population means ± StDev for normalized frequency power. All measures show no differences between the groups despite differences observed at the same age for P6 enucleated animals.

## Discussion

In this study we used bilateral enucleation to examine the role that retina and retinal activity plays in the development of spontaneous (‘background’) activity in the primary visual cortex. Surprisingly, cortical activity recovers within days of bilateral enucleation at P6 and develops nearly normally (Fig1). This occurs despite the fact that spontaneous activity in the retina is responsible for 80-90% of firing in visual cortex and thalamus during the first and second postnatal weeks (Colonnese and Khazipov 2010; Murata and Colonnese 2016). Most importantly, the timing of a major developmental switch in thalamocortical activity (Colonnese and Phillips 2018) is largely unchanged. This switch includes the acquisition of ‘continuous’ background firing, the elimination of development-specific oscillations, and the onset of adult-like frequency distributions indicative of the inhibition-balanced cortical circuit. In this the mammalian cortex appears similar to the zebra fish optic tectum, the network activity patterns remain largely intact following binocular enucleation (Avitan et al. 2017; Pietri et al. 2017). Our data confirm that immature spindle-burst oscillations are an intrinsically generated thalamocortical network oscillation (Luhmann and Khazipov 2018). Under normal circumstances, the duration, occurrence and microstructure of these oscillations arise through an interaction between activity (usually spontaneous) originating in the driving input to thalamus (eg, retina for dLGN; paralimbic cortex for ventral medial thalamus (Hartung et al. 2016)) and the specialized early properties of the thalamocortical circuit (Bitzenhofer et al. 2017). In the absence of the normal driving input, the circuit quickly compensates to generate activity similar to the non-deprived cortex, though at a reduced level (Khazipov et al. 2004). We extend these results by showing that the developmental process that eliminates these early network properties and replaces them with mature sensory processing dynamics--a process that is exquisitely timed to the onset of active sensation (whisking in somatosensory cortex; eye-opening in visual cortex)--is largely independent of the sense organ, at least in the visual system. We identified a number of subtle developments--particularly a delay in the onset of fully continuous background activity and the balance between low-frequency and fast (beta-gamma) activity occurring with eye-opening--that are disrupted by P6 enucleation, but not by P13 enucleation. Thus, changes in retinal activity do not underlie the development of background activity in visual cortex, but the early presence of retina is required to set up circuit conditions required for the exact developmental expression of this activity.

### Thalamocortical homeostatic control of background activity development

Neuronal firing-rates (as well as higher-order structure such as firing-rate correlations and proximity to criticality) are under homeostatic control in more mature juvenile visual cortex, returning to baseline levels within days following eye-lid suture (Hengen et al. 2016; Wu et al. 2020). It is still unclear how and when these homeostatic set points are established. Are spike-rate set-points established by incoming retinal activity levels during early development, or are they an independent function of cortical (or thalamocortical) neurons? Development poses a particularly thorny problem for homeostatic mechanisms as activity is constantly changing. Is this change a result of a changing set-point or is there no set point, with activity determined solely by the inputs? Here we asked whether the developmental increase in firing-rates (Colonnese et al. 2017) and continuity (Shen and Colonnese 2016) is timed by retinal inputs or by changing homeostatic set-points within thalamocortex. Our results suggest the latter. This is remarkable because in the intact system, the macro patterning, as well as upwards of 90% of the synaptic drive to thalamus and cortex, is provided by the retina until late in the second post-natal week when the influence of the retina on cortical activity is reduced (Colonnese et al. 2010; Murata and Colonnese 2016; Gribizis et al. 2019). However retinal input is not the sole determinant of cortical firing rates or patterns. Retinal waves actually provide a net de-synchronizing influence to relay thalamus and cortex (Weliky and Katz 1999; Siegel et al. 2012), with synchronized bursts following rapidly upon removal or silencing of the eyes. The thalamic response to retinal waves is amplified 5-fold by an excitatory feedback loop with visual cortex (Murata and Colonnese 2016). Additional amplification and modulation is likely provided by transient circuits formed between subplate, thalamus and overlying cortex (Kanold and Luhmann 2010) and by inhibitory networks within cortex (Kirmse et al. 2015; Murata and Colonnese 2020). Thus, there is significant potential by thalamic and local cortical circuits to modulate firing rates within homeostatically set ranges.

“True” homeostasis requires the capacity for down-regulation following over activation (Turrigiano 2017), which we did not examine here because of the difficulty in consistently over-activating the developing retina. Thus, it remains unclear whether the consistent shifts in firing-rate and continuity we observed result from a shifting set-point or reflect a changing age-dependent firing-rate maxima that is constantly trying to approach the adult set-point.

In the future it will be important to understand the extent to which, in normal animals, the increase in continuity and firing-rates are driven by shifts in the underlying firing of the retina, which may in fact change quite significantly and contribute to the normal changes observed during development. In fact, matched developmental shifts in firing rate and pattern are likely, with the ultimate output of the system determined by a compromise between retinal, thalamic and cortical homeostatic set points. Evidence for this comes from the subtle changes in silent period duration, continuity and frequency power we observed in the enucleated animals.

### Role of the retina in regulating thalamocortical development

While the developmental trajectory of visual cortex was very similar in P6 enucleated and normal animals, there were some subtle differences in the timing and quality of activity: (1) spindle-burst oscillations did not accelerate between P6 and P10; (2) a second decease in silent-period duration that finalizes the transition to continuous activity was delayed by 2-3 days; (3) background dEEG contained excess low frequency activity that persisted until the end of the study (P16-17). These changes are likely not a result of the loss of the specific temporal patterning provided by retina as (1) retinal waves do not pattern the rapid oscillations within thalamocortex (Colonnese and Khazipov 2010); (2) P13 enucleation did not have any effect on these characteristics. Instead our results suggest that between P6 and P12 retinal absence either prevents circuit maturation or causes circuit adjustments that are required for normal activity after P13 (Fig. 8). One likely locus is for this circuit change is relay thalamus, as the onset of continuous activity originates there and it contributes to the development of dEEG spectral power (Murata and Colonnese 2019). Eyeless mice experience a precocious development of cortical feedback connections, which may contribute to premature reduction of silent periods and to the later developmental delay (Seabrook et al. 2013). The reorganization of subplate and interneuron connectivity which occurs in the second post-natal week is another potential locus for aberrant circuit development following P6 enucleation (Lim et al. 2018; Kanold et al. 2019). One process unlikely to contribute is sprouting of other sensory inputs into visual thalamus or thalamic regions to visual cortex, as P6 is after the period when calcium waves establish cortical sensory identity (Moreno-Juan et al. 2017).

**Figure 8.**
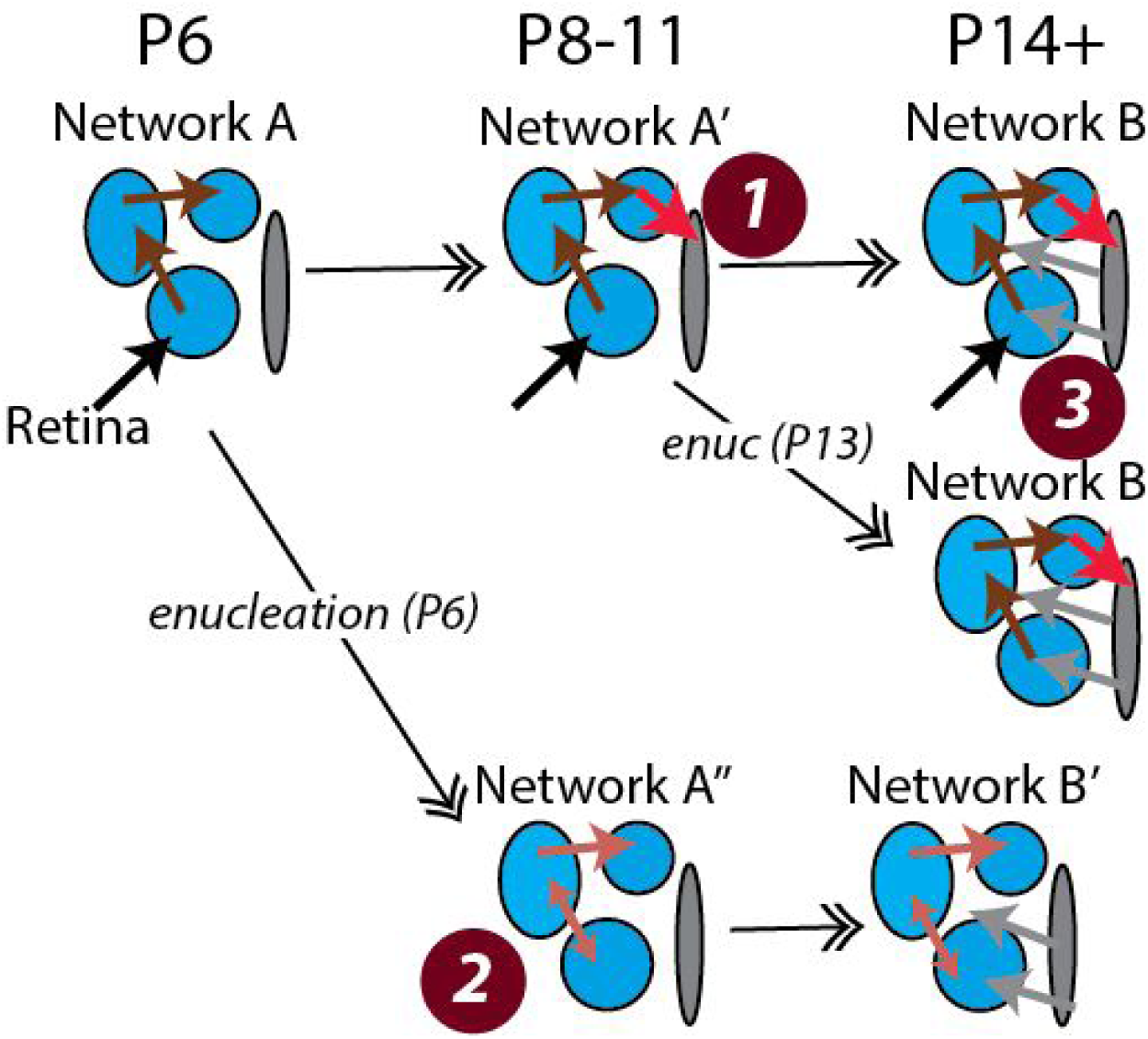
Schematic of potential circuit basis of enucleation effects. Normal development (top row) consists of minor changes (Network A to A’) in circuits until late in the second week when the incorporation of a novel input 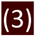) shifts the network (Network B) to continuous activity. Part of the shift in from Network A to A’ includes development of connection 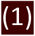 that has little role until the onset of Network B. P6 enucleation (bottom row) results in a slightly modified early network (A’’) that includes synaptic modifications that maintains spike-rates in the absence of the retina 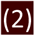. Later incorporation of gray inputs occurs normally, resulting in a modified mature network (B’). Late enucleation occurs after development of 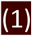 thus incorporation of gray input results in normal network B.

### Clinical relevance

Continuous EEG monitoring has been formally recommended for use in all neonates at risk of brain injury by the American Clinical Neurophysiology Society (Shellhaas et al. 2011). The EEG’s utility is based on the well described, age-linked, changes in the continuity of background activity as well as the stereotyped evolution of spontaneous activity patterns, which are also observed in animal models including mice (Cirelli and Tononi 2015). The relay thalamus appears to be the central organizer of this development, at least in primary sensory cortex (Murata and Colonnese 2019). However, whether thalamic activity changes simply reflect changing input or result from maturing thalamic circuits has been unclear. Our data clearly support the latter hypothesis, as removal of the eyes did not significantly alter the timing or form of activity development. Our data would predict that pre thalamic lesions to the primary ‘driver’ input would be difficult to detect by clinical EEG, and further suggest that thalamic lesions should be suspected when there is significant developmental delay or discontinuity in the cortical EEG of infants.

## Acknowledgements

This work was supported by R01EY022730 and R01NS106244.

